# Differential relationships between habitat fragmentation and within-population genetic diversity of three forest-dwelling birds

**DOI:** 10.1101/004903

**Authors:** Benjamin Zuckerberg, Matthew D. Carling, Roi Dor, Elise D. Ferree, Garth M. Spellman, Andrea K. Townsend

## Abstract

Habitat fragmentation is a major driver of environmental change affecting wildlife populations across multiple levels of biological diversity. Much of the recent research in landscape genetics has focused on quantifying the influence of fragmentation on genetic variation among populations, but questions remain as to how habitat loss and configuration influences within-population genetic diversity. Habitat loss and fragmentation might lead to decreases in genetic diversity within populations, which might have implications for population persistence over multiple generations. We used genetic data collected from populations of three species occupying forested landscapes across a broad geographic region: Mountain Chickadee (*Poecile gambeli*; 22 populations), White-breasted Nuthatch (*Sitta carolinensis*; 13 populations) and Pygmy Nuthatch (*Sitta pygmaea*; 19 populations) to quantify patterns of haplotype and nucleotide diversity across a range of forest fragmentation. We predicted that fragmentation effects on genetic diversity would vary depending on dispersal capabilities and habitat specificity of the species. Forest aggregation and the variability in forest patch area were the two strongest landscape predictors of genetic diversity. We found higher haplotype diversity in populations of *P. gambeli* and *S. carolinensis* inhabiting landscapes characterized by lower levels of forest fragmentation. Conversely, *S. pygmaea* demonstrated the opposite pattern of higher genetic diversity in fragmented landscapes. For two of the three species, we found support for the prediction that highly fragmented landscapes sustain genetically less diverse populations. We suggest, however, that future studies should focus on species of varying life-history traits inhabiting independent landscapes to better understand how habitat fragmentation influences within-population genetic diversity.

## INTRODUCTION

Ecologists often study how habitat composition and configuration influences the abundance and occurrence of organisms (Turner et al. 2001, Fahrig 2003), but in recent years, the field of landscape genetics has focused on how these landscape features influence genetic variation (Anderson et al. 2010, Sork and Waits 2010, Storfer et al. 2010). Landscape genetics studies have successfully incorporated the role of isolation and geographic distance in explaining genetic variation among populations (e.g., Jenkins et al. 2010); however, there has been a persistent interest in examining how habitat composition and configuration influences genetic diversity within populations (Bruggeman et al. 2010, Sork and Waits 2010, Thomassen et al. 2010). This interest has been driven, in part, by the concern that habitat fragmentation might lead to decreases in genetic diversity within animal populations, which might have strong implications for the persistence of populations over multiple generations (Frankham et al. 2010).

The process of habitat loss and fragmentation produces landscapes characterized by isolated habitat patches of decreasing size and continuity and has important implications on genetic diversity. Landscapes characterized by habitat loss and fragmentation are thought to support smaller effective population sizes (Wilcove 1985, Debinski and Holt 2000, Chalfoun et al. 2002), which neutral theory predicts will lead to decreased genetic diversity (Hartl and Clark 2007). Empirical evidence seems to support this prediction as birds with relatively larger populations and broader distributions are genetically more diverse (Moller et al. 2008). For many species, one of the mechanisms driving this pattern is that fragmentation causes decreased rates of individual movement and dispersal among isolated habitat patches (Belisle et al. 2001), leading to a subdivided and more variable population (Boulinier et al. 1998, Boulinier et al. 2001, Donovan and Flather 2002). In turn, random genetic drift and reduced gene flow leads to an eventual decline of within-population genetic diversity as a result of declining and/or disconnected populations (Templeton et al. 1990, Young et al. 1996, Keyghobadi 2007).

Comparative studies on the genetic impact of habitat fragmentation among populations generally support the prediction of reduced genetic diversity in fragmented landscapes (Keyghobadi 2007). Many of these studies, however, are plagued by analytical and methodological limitations. First, studies of habitat fragmentation often confound habitat loss with the process of habitat fragmentation per se (the spatial breaking apart of habitat independent of habitat loss) (Fahrig 2003). A potential reason for this limitation results from a lack of replication of independent landscapes by which to assess variability in habitat configuration (McGarigal and Cushman 2002). Second, most comparative studies compare genetic structure within a control and fragmented landscape in the same geographic region. As a result, it is difficult to identify which aspects of fragmentation might be driving changes in genetic responses and differentiate those from broader-scaled regional patterns (Johansson et al. 2005). Third, few studies focus on multiple species with varying life history characteristics. It is likely that species of different characteristics of dispersal and habitat specificity could mediate the genetic response to fragmentation at a landscape scale (Van Houtan et al. 2007, Bowie 2011, Callens et al. 2011). Genetic diversity is a population characteristic that should be particularly sensitive to the spatial distribution of habitat, and there is a need to evaluate the characteristics of habitat loss and fragmentation that could lead to genetic variation among multiple species inhabiting a continuum of fragmentation.

We quantified patterns of within-population genetic diversity for bird populations exposed to a range of habitat fragmentation for three species of passerine birds found throughout the western United States: Mountain Chickadee (*Poecile gambeli*), White-breasted Nuthatch (*Sitta carolinensis*) and Pygmy Nuthatch (*Sitta pygmaea*). Previous molecular analyses have suggested that some of the range-wide patterns of genetic structure and among-population diversity in these three species can be explained by historical isolation and subsequent expansion from glacial refugia in the Pleistocene. In *P. gambeli*, two clades are consistent with isolation in two refugia during glacial advances (Spellman et al. 2007), and among western populations of *S. carolinensis*, two of three clades decrease in nucleotide diversity with increasing latitude, a pattern consistent with expansion from southern glacial refugia (Spellman and Klicka 2007). For *S. pygmaea*, expansion from a single glacial refugium in southern coastal California may have led to a positive correlation of nucleotide diversity with longitude (Spellman and Klicka 2006). However, these studies primarily focused on phylogeographic patterns of genetic diversity among populations as opposed to genetic characteristics within independent populations.

Our objective was to use existing genetic data to test the relationship between habitat fragmentation and within-population genetic diversity for the above three forest-breeding bird species across a range of fragmentation. We predicted that the relationship between fragmentation and within-population genetic diversity, as measured by haplotype and nucleotide diversity, would vary among species according to their dispersal capabilities and habitat requirements. Specifically, we predicted that the genetic diversity of the *P. gambeli*, with its intermediate dispersal distance (McCallum et al. 1999) and habitat specificity might not vary with fragmentation. We predicted that genetic diversity would be higher for the *S. carolinensis* in fragmented landscapes, because of their longer dispersal distances (Grubb and Pravosudov 2008) and relatively diverse habitat associations. Finally, we predicted that genetic diversity would decrease in fragmented landscapes for the *S. pygmaea*, with its relatively poor dispersal capabilities (Kingery and Ghalambor 2001) and strict habitat associations, because it would be less likely to disperse across a fragmented landscape (Van Houtan et al. 2007). Generalist species that exhibit relatively greater rates of dispersal should be less susceptible to the impacts of habitat fragmentation and its associated processes that erode local genetic diversity over time (Mech and Hallet 2001, Vandergast et al. 2007, Bowie 2011).

## METHODS

### Study species and genetic sampling

The distribution of each of our study species, *P. gambeli, S. carolinensis*, and *S. pygmaea*, matches the distribution of their preferred forest habitats, coniferous forests, pine-oak and long-needle pine, respectively. *Poecile gambeli* generally prefers montane coniferous forests and areas dominated by pine, spruce-fir (Picea-Abies), and piñon-juniper (Pinus-Juniperus) (McCallum et al. 1999). They will occur in mixed coniferous-deciduous habitats, but show preferential use of conifer stands in these settings (McCallum et al. 1999). *Poecile gambeli* generally does not migrate, but most yearlings exhibit natal dispersal, although young dispersers sometimes move only as far as adjacent family groups (McCallum et al. 1999). *Sitta carolinensis* is more closely associated with mature deciduous woodland, but can also breed in mixed deciduous, coniferous forest, and a variety of montane woodland habitats (Grubb and Pravosudov 2008). They exhibit somewhat irruptive migration and disperse up to 10 km in their first year, and although little is known about natal dispersal distances for *S. carolinensis* (Grubb and Pravosudov 2008), mean natal dispersal distances were less than 10 km in both a Belgian population (Matthysen and Schmidt 1987) and German population (Winkel 1989) of Eurasian nuthatches (*Sitta europaea*). Finally, *S. pygmaea* is the most specific in terms of habitat requirements. *Sitta pygmaea* demonstrates a strong and almost exclusive preference for long-needled pine forests and are co-extensive with that of ponderosa pine (*Pinus ponderosa*), Jeffrey pine (*Pinus jeffreyi*), and similar species (Kingery and Ghalambor 2001). They are also the most sedentary of the three study species, with no migration and very short natal dispersal distances (< 500 m; Kingery and Ghalambor 2001). Recorded mean natal dispersal distance for *S. pygmaea* centers around 287 m (range = 0-533 m; Norris 1958). Evidence from individual populations also suggests that each species is most likely to occur in relatively intact forested areas when compared to fragmented landscapes (Brawn and Balda 1988, McCallum et al. 1999, Villard et al. 1999). Because of their wide distribution across western North America and reliance on forest habitats, these species created a suitable system to examine the influence of habitat fragmentation on within-population genetic diversity.

For each species, we used available sequences of the mitochondrial gene NADH dehydrogenase subunit 2 (ND2; 1041 bp) from previously published phylogeographic studies on the three species (Spellman and Klicka 2006, 2007, Spellman et al. 2007). We used previously generated ND2 sequences from 216 *P. gambeli* individuals (22 populations), 92 *S. carolinensis* individuals (13 populations) and 132 *S. pygmaea* individuals (19 populations; Fig. 1; Table 1). The location of sampling sites emphasized protected areas with limited anthropogenic disturbance (e.g., National Forests, State Parks, and State Game Preserves).

**FIGURE 1.**
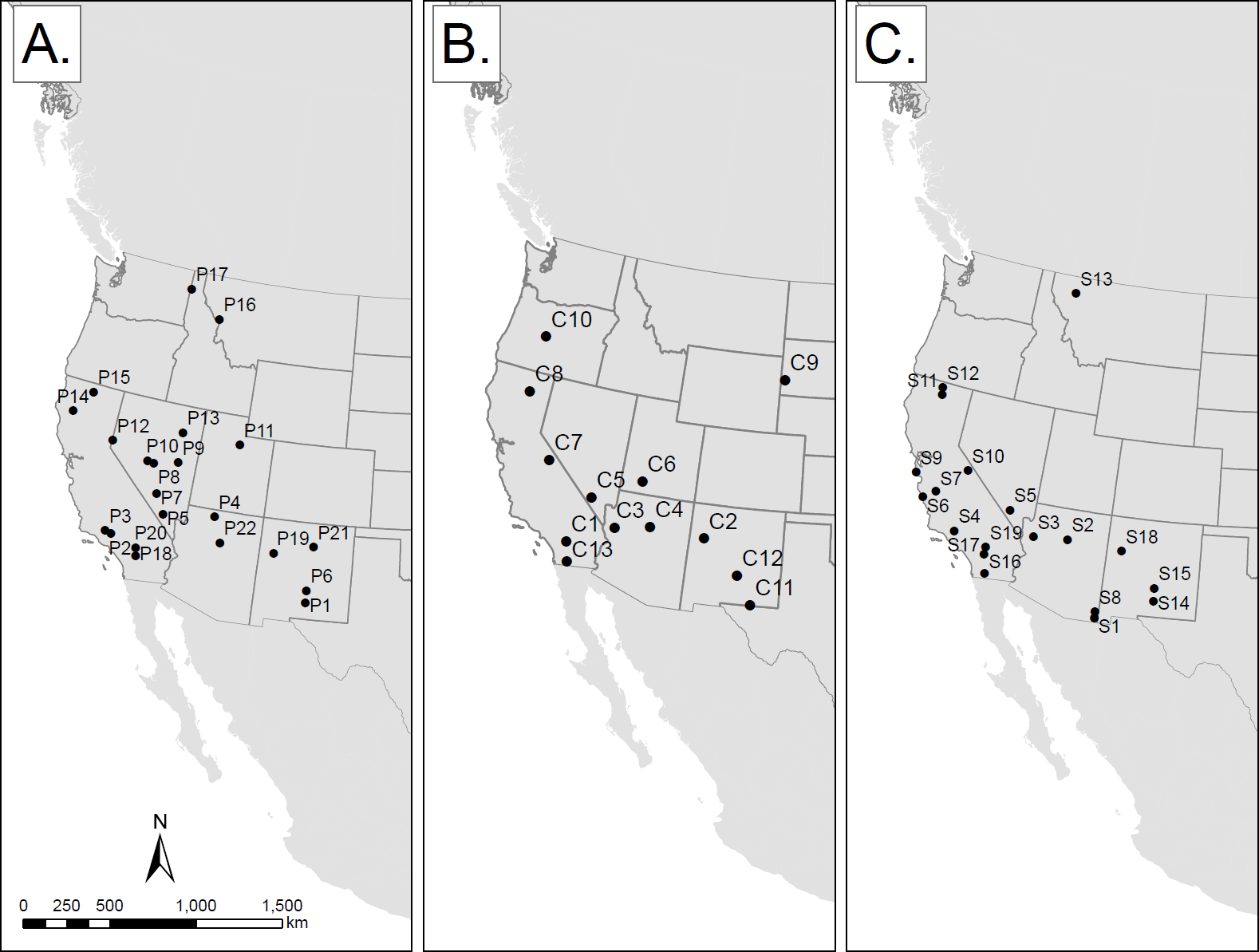
Geographic locations of sampling points for *Poecile gambeli* (A.), *Sitta carolinensis* (B.), and *Sitta pygmaea* (C.) in western USA. Labels represent population identifiers (see Table 1).

**TABLE 1.**
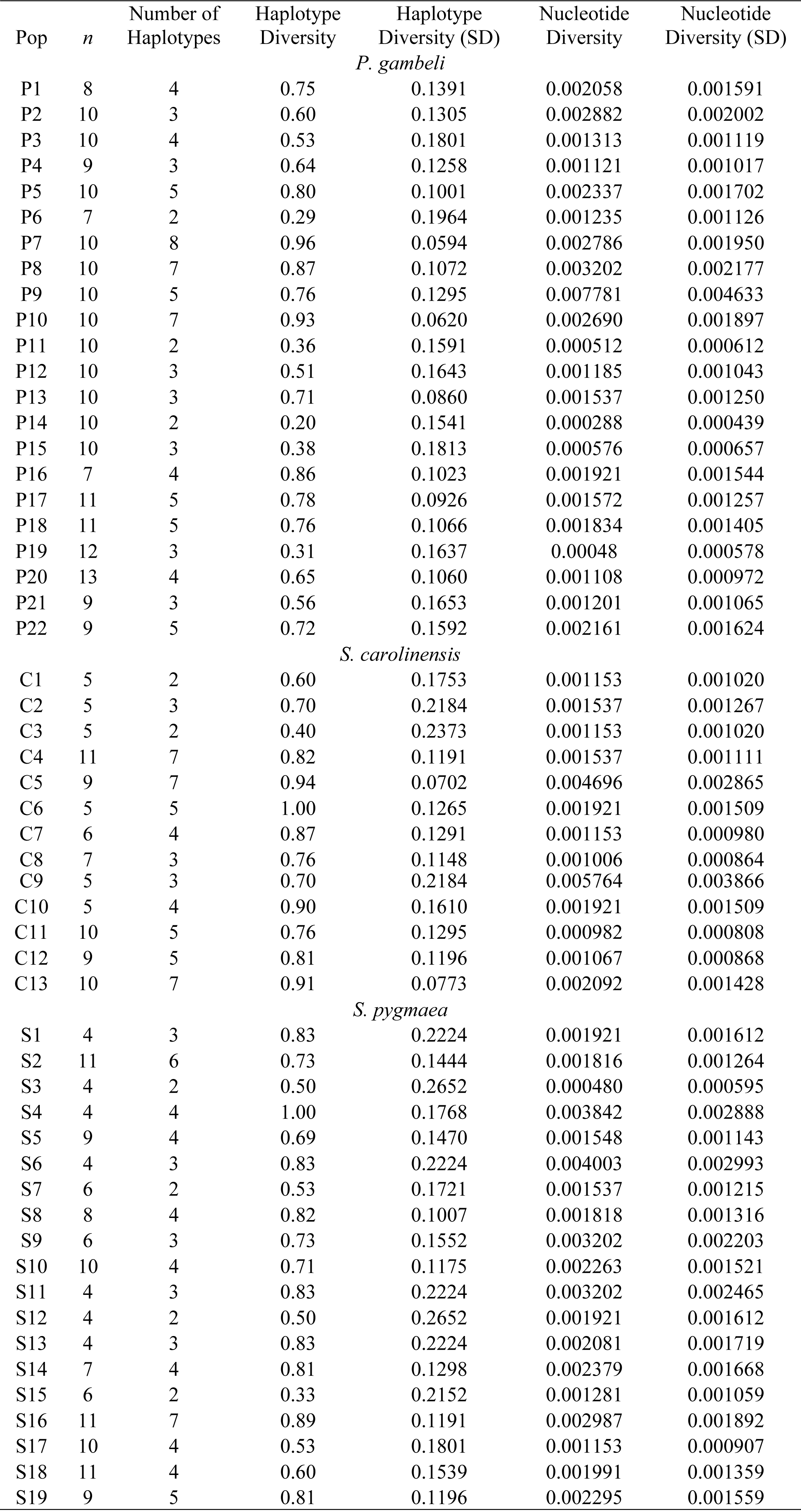
Population identifier, number of individuals sampled (*n*), haplotype number (number of unique haplotypes in each population), haplotype diversity and standard deviation, and pairwise nucleotide diversity and standard deviation (SD), derived from complete sequences of the mitochondrial gene NADH dehydrogenase subunit 2 (ND2), for each population from the three species used in this study.

| Pop | $n$ | Number of Haplotypes | Haplotype Diversity | Haplotype Diversity (SD) | Nucleotide Diversity | Nucleotide Diversity (SD) |
| --- | --- | --- | --- | --- | --- | --- |
| *P. gambeli* |  |  |  |  |  |  |
| P1 | 8 | 4 | 0.75 | 0.1391 | 0.002058 | 0.001591 |
| P2 | 10 | 3 | 0.60 | 0.1305 | 0.002882 | 0.002002 |
| P3 | 10 | 4 | 0.53 | 0.1801 | 0.001313 | 0.001119 |
| P4 | 9 | 3 | 0.64 | 0.1258 | 0.001121 | 0.001017 |
| P5 | 10 | 5 | 0.80 | 0.1001 | 0.002337 | 0.001702 |
| P6 | 7 | 2 | 0.29 | 0.1964 | 0.001235 | 0.001126 |
| P7 | 10 | 8 | 0.96 | 0.0594 | 0.002786 | 0.001950 |
| P8 | 10 | 7 | 0.87 | 0.1072 | 0.003202 | 0.002177 |
| P9 | 10 | 5 | 0.76 | 0.1295 | 0.007781 | 0.004633 |
| P10 | 10 | 7 | 0.93 | 0.0620 | 0.002690 | 0.001897 |
| P11 | 10 | 2 | 0.36 | 0.1591 | 0.000512 | 0.000612 |
| P12 | 10 | 3 | 0.51 | 0.1643 | 0.001185 | 0.001043 |
| P13 | 10 | 3 | 0.71 | 0.0860 | 0.001537 | 0.001250 |
| P14 | 10 | 2 | 0.20 | 0.1541 | 0.000288 | 0.000439 |
| P15 | 10 | 3 | 0.38 | 0.1813 | 0.000576 | 0.000657 |
| P16 | 7 | 4 | 0.86 | 0.1023 | 0.001921 | 0.001544 |
| P17 | 11 | 5 | 0.78 | 0.0926 | 0.001572 | 0.001257 |
| P18 | 11 | 5 | 0.76 | 0.1066 | 0.001834 | 0.001405 |
| P19 | 12 | 3 | 0.31 | 0.1637 | 0.00048 | 0.000578 |
| P20 | 13 | 4 | 0.65 | 0.1060 | 0.001108 | 0.000972 |
| P21 | 9 | 3 | 0.56 | 0.1653 | 0.001201 | 0.001065 |
| P22 | 9 | 5 | 0.72 | 0.1592 | 0.002161 | 0.001624 |
| *S. carolinensis* |  |  |  |  |  |  |
| C1 | 5 | 2 | 0.60 | 0.1753 | 0.001153 | 0.001020 |
| C2 | 5 | 3 | 0.70 | 0.2184 | 0.001537 | 0.001267 |
| C3 | 5 | 2 | 0.40 | 0.2373 | 0.001153 | 0.001020 |
| C4 | 11 | 7 | 0.82 | 0.1191 | 0.001537 | 0.001111 |
| C5 | 9 | 7 | 0.94 | 0.0702 | 0.004696 | 0.002865 |
| C6 | 5 | 5 | 1.00 | 0.1265 | 0.001921 | 0.001509 |
| C7 | 6 | 4 | 0.87 | 0.1291 | 0.001153 | 0.000980 |
| C8 | 7 | 3 | 0.76 | 0.1148 | 0.001006 | 0.000864 |
| C9 | 5 | 3 | 0.70 | 0.2184 | 0.005764 | 0.003866 |
| C10 | 5 | 4 | 0.90 | 0.1610 | 0.001921 | 0.001509 |
| C11 | 10 | 5 | 0.76 | 0.1295 | 0.000982 | 0.000808 |
| C12 | 9 | 5 | 0.81 | 0.1196 | 0.001067 | 0.000868 |
| C13 | 10 | 7 | 0.91 | 0.0773 | 0.002092 | 0.001428 |
| *S. pygmaea* |  |  |  |  |  |  |
| S1 | 4 | 3 | 0.83 | 0.2224 | 0.001921 | 0.001612 |
| S2 | 11 | 6 | 0.73 | 0.1444 | 0.001816 | 0.001264 |
| S3 | 4 | 2 | 0.50 | 0.2652 | 0.000480 | 0.000595 |
| S4 | 4 | 4 | 1.00 | 0.1768 | 0.003842 | 0.002888 |
| S5 | 9 | 4 | 0.69 | 0.1470 | 0.001548 | 0.001143 |
| S6 | 4 | 3 | 0.83 | 0.2224 | 0.004003 | 0.002993 |
| S7 | 6 | 2 | 0.53 | 0.1721 | 0.001537 | 0.001215 |
| S8 | 8 | 4 | 0.82 | 0.1007 | 0.001818 | 0.001316 |
| S9 | 6 | 3 | 0.73 | 0.1552 | 0.003202 | 0.002203 |
| S10 | 10 | 4 | 0.71 | 0.1175 | 0.002263 | 0.001521 |
| S11 | 4 | 3 | 0.83 | 0.2224 | 0.003202 | 0.002465 |
| S12 | 4 | 2 | 0.50 | 0.2652 | 0.001921 | 0.001612 |
| S13 | 4 | 3 | 0.83 | 0.2224 | 0.002081 | 0.001719 |
| S14 | 7 | 4 | 0.81 | 0.1298 | 0.002379 | 0.001668 |
| S15 | 6 | 2 | 0.33 | 0.2152 | 0.001281 | 0.001059 |
| S16 | 11 | 7 | 0.89 | 0.1191 | 0.002987 | 0.001892 |
| S17 | 10 | 4 | 0.53 | 0.1801 | 0.001153 | 0.000907 |
| S18 | 11 | 4 | 0.60 | 0.1539 | 0.001991 | 0.001359 |
| S19 | 9 | 5 | 0.81 | 0.1196 | 0.002295 | 0.001559 |

### Genetic data analysis

Using ARLEQUIN v3.11, we estimated two indices of within-population genetic diversity: haplotype diversity and pairwise nucleotide diversity (Excoffier et al. 2005). Haplotype diversity is the probability that two randomly chosen haplotypes are different in the sample, whereas nucleotide diversity is the average pairwise nucleotide difference between two sequences in each population. Nucleotide diversity is haplotype diversity at the nucleotide level; therefore, it captures not only how many different haplotypes are in a population, but also how different those haplotypes are from each other. Low haplotype and nucleotide diversity can indicate low underlying effective population size (Avise 2000), the influence of recent expansions from glacial refugia (Spellman and Klicka 2006) or cyclical range shifts (Jaarola et al. 1999). These two indices capture different aspects of genetic variation, but they may be statistically correlated. We performed separate analyses for each of the species including sequences from all available individuals from each population (Table 1).

### Landscape analysis

We defined landscapes as a 10 km buffer around each sampled population’s centroid to represent the spatial extent encountered by most dispersing individuals within a single generation. Although natal dispersal distances have not been fully characterized for our focal species, available evidence suggests that these maximum dispersal distances are within a 10 km-radius landscape (see *Study species and genetic sampling*).

We analyzed the 2001 National Land Cover Data (NLCD) to characterize the cover and configuration of forest cover in the 10 km-radius landscape. The 2001 NLCD consists of 16 land cover classes modeled over the conterminous United States at a 30 m cell resolution and a 0.40 ha minimum mapping unit (Homer et al. 2007). We calculated the proportion of each land cover type within each landscape, but because the landscapes tended to be dominated by two primary cover types (deciduous or coniferous forest and early-successional cover types), we chose to aggregate the land cover classes to represent a binary habitat cover map of forest (including deciduous/mixed forest or evergreen forest types) and non-forest (including early-successional) cover types. Although there are many metrics available for quantifying landscape heterogeneity (Cushman et al. 2008), we quantified fragmentation using a subset of metrics that have been shown to characterize aspects of forest fragmentation important for forest-breeding birds (e.g., forest patch size variability and forest aggregation) (Table 2). The initial set of metrics used to describe forest fragmentation included the percentage of landscape classified as forest (Forest), area-weighted mean forest patch area (AREA_AM), forest patch area coefficient of variation (AREA_CV), mean nearest neighbor distance (ENN), nearest neighbor distance coefficient of variation (ENN_CV), area-weighted mean edge contrast (ECON), similarity index (SIMI), and clumpiness index (CLUMPY) (McGarigal et al. 2002). Multicollinearity is a common problem when using fragmentation metrics as predictors, and for each species we examined scatter plots to identify the least correlated (*r* < 0.5) set of predictor variables. We included the proportion of forest cover at a 100 km radius landscape (FOR) as a measure of regional habitat availability and to account for processes that might be occurring over broader spatial scales and maximum dispersal events (Tittler et al. 2009). We calculated all landscape metrics in FRAGSTATS (McGarigal et al. 2002).

**TABLE 2.**
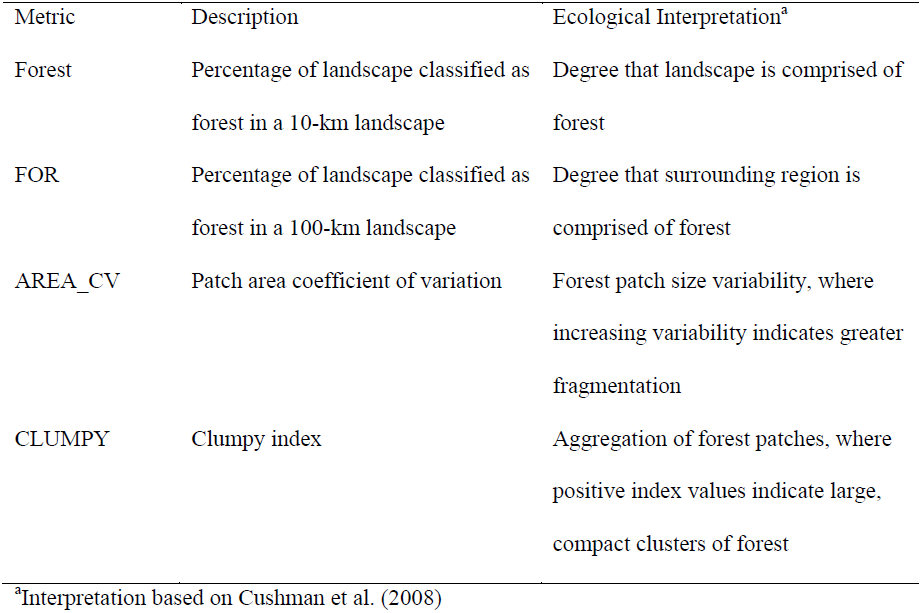
Metrics used to quantify forest fragmentation within 10 km radius-landscapes. We include interpretations for only those metrics retained in the top models identified through model selection.

| Metric | Description | Ecological Interpretation<sup>a</sup> |
| --- | --- | --- |
| Forest | Percentage of landscape classified as forest in a 10-km landscape | Degree that landscape is comprised of forest |
| FOR | Percentage of landscape classified as forest in a 100-km landscape | Degree that surrounding region is comprised of forest |
| AREA_CV | Patch area coefficient of variation | Forest patch size variability, where increasing variability indicates greater fragmentation |
| CLUMPY | Clumpy index | Aggregation of forest patches, where positive index values indicate large, compact clusters of forest |

<sup>a</sup>Interpretation based on Cushman et al. (2008)

### Statistical analysis

For each species, we used a generalized linear model with a Gaussian error distribution (GLM) (Faraway 2006) and AIC*_c_*-based model selection (Burnham and Anderson 2002) to identify the most parsimonious model of the relationship between within-population genetic diversity and forest fragmentation. Forest cover (at the 10 km landscape-scale), and regional forest availability (at the 100 km landscape-scale) was included in every model to statistically examine the variation in forest configuration while controlling for forest composition. All fragmentation metrics were standardized for analysis. As the number of individuals sampled in a population can potentially influence genetic measures (Landguth et al. 2010), we ran two-sided Grubb’s tests for each species to test for the presence of outliers in the number of sampled individuals (Grubbs 1950). A Grubb’s test detects one outlier at a time, and the outliers are expunged from the dataset and the test is iterated until no outliers are detected. Following the identification and elimination of outliers, sample size (*n*) was included as a covariate in model selection. For all model runs, we included a model that included only forest cover (at both landscape and regional scales). We calculated the number of model parameters (*K*), AIC*_c_* values, ΔAIC*_c_*, and Akaike weights (*w_i_*). Models with a ΔAIC*_c_* < 2 were considered equivalent (Burnham and Anderson 2002). We calculated Nagelkerke R^2^ for assessing the explained variation of the global model in a likelihood framework (Nagelkerke 1991). We performed model selection using R (Version 2.10.1; R Development Core Team 2009) with the extension package MuMIn (Barton 2010).

### Spatial analysis

We checked for spatial autocorrelation to quantify whether patterns in within-population genetic diversity were associated with broader latitudinal gradients fragmentation (Spellman and Klicka 2006, 2007, Spellman et al. 2007) after accounting for the variation explained by forest cover and fragmentation. We calculated spatial correlograms (Moran’s *I*) on residuals of the model with strongest support through model selection (Model 1 for each species) at different lag distances (maximum lag distance = 200 km) and tested whether spatial autocorrelation was significant through resampling with 10,000 iterations (alpha < 0.05) (Dormann et al. 2007). We performed all spatial analyses using the R extension package ncdf (Dormann et al. 2007).

## RESULTS

### Genetic diversity within populations

We found within-population genetic diversity to be variable among populations (within-species) both for haplotype diversity (*P. gambeli*: mean = 0.62 ± 0.26; *S. carolinensis*: mean = 0.77 ± 0.24; *S. pygmaea*: mean = 0.67 ± 0.23) and for pairwise nucleotide diversity (*P. gambeli*: mean = 0.0019 ± 0.0015; *S. carolinensis*: mean = 0.0021 ± 0.0015; *S. pygmaea*: mean = 0.0021 ± 0.0010). As expected, haplotype diversity was correlated with nucleotide diversity for all three species (*P. gambeli*: *r*^2^ = 0.41, *P* < 0.001; *S. carolinensis*: *r*^2^ = 0.22, *P* = 0.04; *S. pygmaea*: *r*^2^ = 0.63, *P* < 0.001).

### Patterns of forest fragmentation

We found that the local 10 km landscapes surrounding the sampling locations consisted of two primary land cover types, forested upland (characterized by natural or semi-natural wood vegetation greater than 6 m tall) and shrubland habitats (characterized by natural or semi-natural wood vegetation with aerial stems less than 6 m tall), and supported the analysis of these sites as binary (forest vs. non-forest) landscapes. Shrubland (non-forest habitats) is a classification representing natural vegetation (non-agricultural) as areas dominated by shrubs less than 5 m tall, and is often also associated with grasses, sedges, herbs, and non-vascular vegetation.

For *P. gambeli* populations we found forest composition (10 km radius) ranging from 25% to 95% (mean = 62%) and regional forest composition (100 km) ranging from 0% to 70% (mean = 21%). We found similar patterns for the landscapes surrounding *S. carolinensis* populations, we found that forest composition ranged from 23% to 87% (mean = 61%) and regional forest composition ranged from 1% to 50% (mean = 21%). *Sitta pygmaea* landscapes were also characterized by a combination of forest and shrubland cover types with forest composition ranging from 2% to 96% (mean = 50%) and regional forest composition from 0% to 65% (mean = 14%).

### Genetic diversity and forest fragmentation

Normal-probability plots and histograms showed that normality for regression-model residuals was met. For *P. gambeli*, sample size ranged from 4-19 individuals and the outlier test was significant (U = 0.40, *P* = 0.003). As such, we iteratively removed populations with fewer than 5 and greater than 14 individuals sampled. Re-running the outlier test demonstrated that no more outliers were detected. The uncorrelated set of metrics included AREA_CV and CLUMPY that resulted in eight separate models. For haplotype diversity, we found that two out of four models had support (ΔAIC*_c_* < 2; Table 3). The model with the strongest support (*w_i_* = 0.46) included the forest cover only model, while the second best model (*w_i_* = 0.18) included only CLUMPY. For *P. gambeli*, haplotype diversity was slightly lower in landscapes with higher forest cover at landscape (Forest; *β* = – 0.13 ± 0.22) and regional scales (FOR; *β* = – 0.03 ± 0.23), but these relationships were highly variable. When forest aggregation (CLUMPY) was included in the model (Model 1; Table 3), we found that haplotype diversity for *P. gambeli* was higher in landscapes with a higher degree of forest aggregation (CLUMPY; *β* = 0.06 ± 0.05; Fig. 2A). The Nagelkerke R^2^ for the global model was 0.41. For nucleotide diversity, we found that one model had the strongest support (*w_i_* = 0.60). Similar to haplotype diversity, we found that nucleotide diversity was higher in landscapes of higher forest aggregation (CLUMPY; *β* = 0.79×10^−3^ ± 0.29×10^−,3^; Fig. 2B).

**FIGURE 2.**
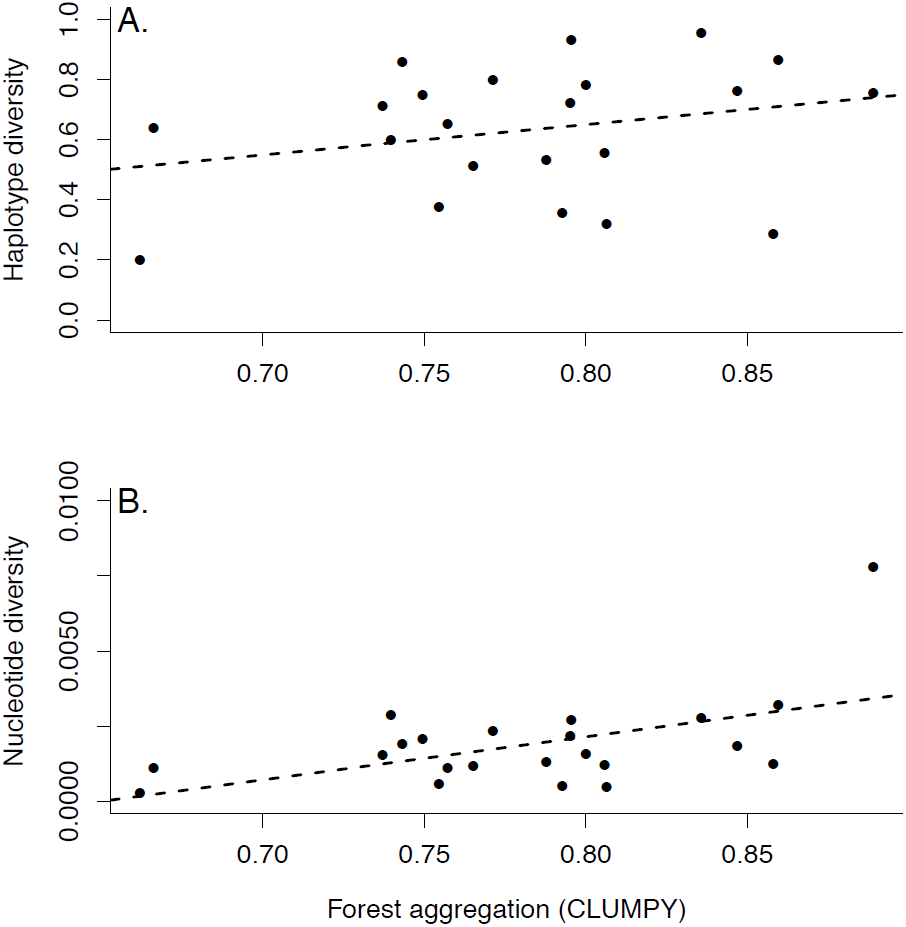
For *P. gambeli*, haplotype diversity (A.) and nucleotide diversity (B.) was higher in landscapes characterized by a higher degree of forest aggregation (CLUMPY).

**TABLE 3.**
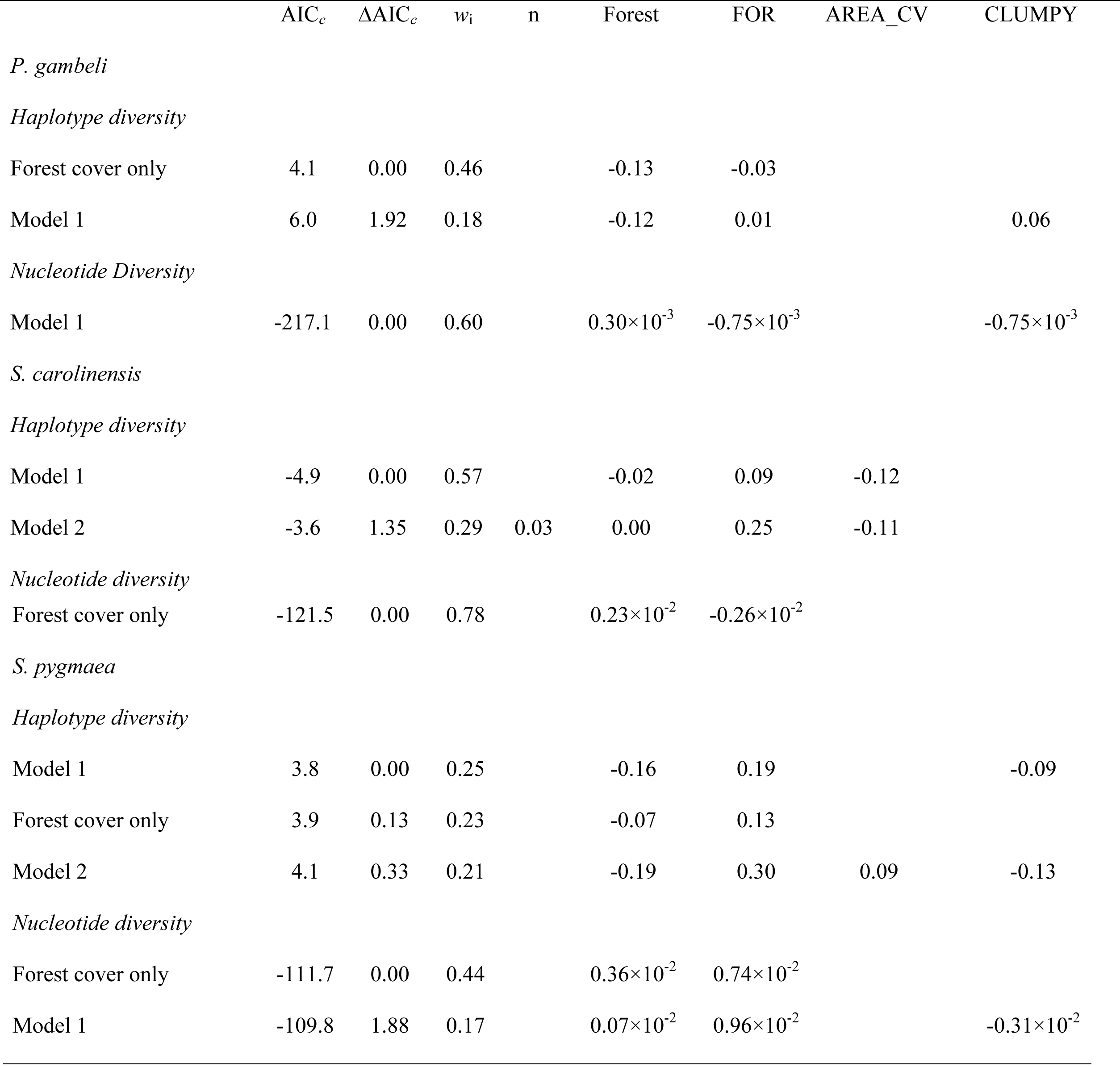
Model selection results testing the relationship between fragmentation and genetic diversity for *P. gambeli*, *S. carolinensis* and *S. pygmaea*. We report Akaike’s second-order information criterion (AIC*_c_*), Akaike weights (*w*_i_) and associated parameter estimates for the fragmentation metrics included in each model. Only models with strong support (ΔAIC*_c_* < 2) are presented. Forest composition at the 10 km (Forest) and 100 km (FOR) landscape scales.

For *S. carolinensis*, sample size ranged from 5-11 individuals, no outliers were detected (U = 0.70, *P* = 1), and the least correlated set of fragmentation metrics for *S. carolinensis* were AREA_CV, SIMI, and CLUMPY. Combinations of these variables resulted in eight separate models. For haplotype diversity, we found that two models had the highest support in model selection (*w_i_* = 0.86). The top model (*w_i_* = 0.57) only included forest patch size variability as a fragmentation metric (AREA_CV; Table 3). We found that *S. carolinensis* populations inhabiting more fragmented landscapes with a higher variability of forest patch sizes (high AREA_CV) had lower haplotype diversity (*β* = – 0.12 ± 0.03; Fig. 3). Haplotype diversity was also slightly higher in more forested landscapes at regional scales (*β* = 0.09 ± 0.23), although this was highly variable. The second most supported model (*w_i_* = 0.29), included sample size (*n*) and showed a slightly positive relationship with haplotype diversity (*β* = 0.03 ± 0.01). The Nagelkerke R^2^ for the global model was 0.78. When analyzing nucleotide diversity the model including only forest cover had the most support (*w_i_* = 0.78; Table 3) suggesting that none of the fragmentation metrics provided sufficient explanatory power in describing the variation in nucleotide diversity for *S. carolinensis*. In these models, nucleotide diversity was higher in more forested landscapes at a local scale, but slightly lower in forested regions.

**FIGURE 3.**
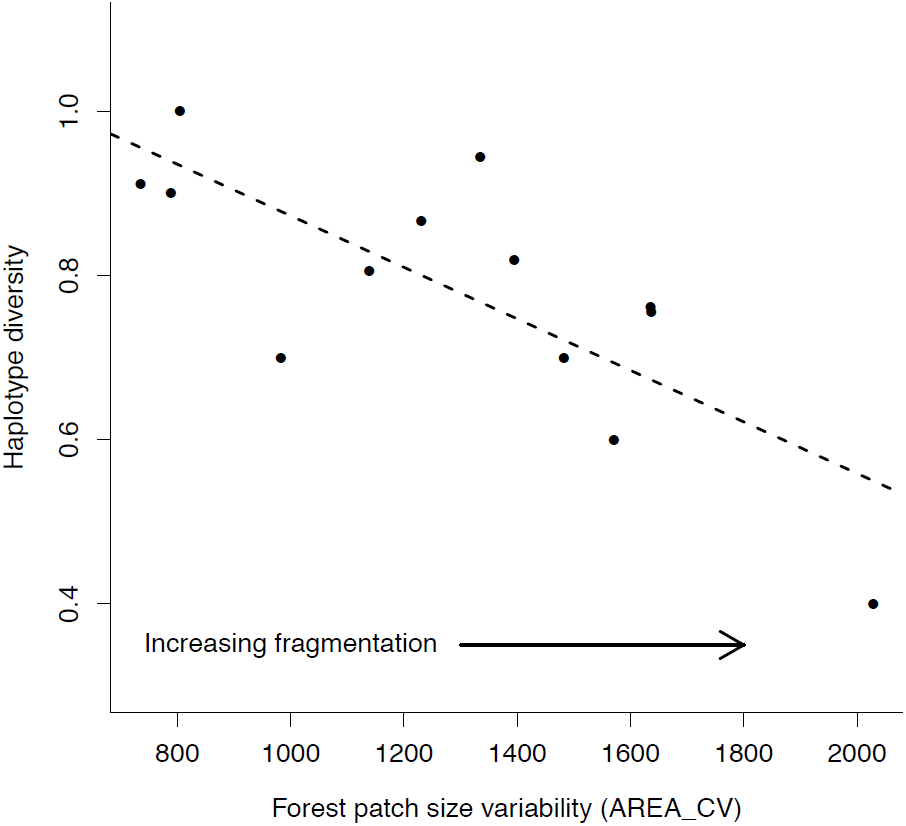
For *S. carolinensis* populations, haplotype diversity was lower in more fragmented landscapes characterized by higher variation in forest patch size (AREA_CV).

*Sitta pygmaea* populations ranged from 4-11 individuals sampled and the outlier test was not significant (U = 0.82, *P* = 1). We included three uncorrelated fragmentation metrics including AREA_CV, ENN_CV, and CLUMPY in model selection. Three out of six models had support (Δ AIC*_c_* < 2), and the final model set included the model with forest cover only (Table 3). In contrast to *P. gambeli* and *S. carolinensis*, we found that the haplotype diversity of *S. pygmaea* populations generally increased in more fragmented landscapes (Nagelkerke R^2^ = 0.47). Haplotype diversity was lower in landscapes with higher forest aggregation (CLUMPY; *β* = −0.09 ± 0.04; Fig. 4) and higher in landscapes with more forest patch size variability (AREA_CV; *β* = 0.10 ± 0.04). Nucleotide diversity showed the same general patterns including the forest cover only with highest support, given the data (*w_i_* = 0.44), and lower nucleotide diversity in landscape characterized by higher forest aggregation (CLUMPY; *β* = −0.31×10^−2^ ± 0.02×10^−2^).

**FIGURE 4.**
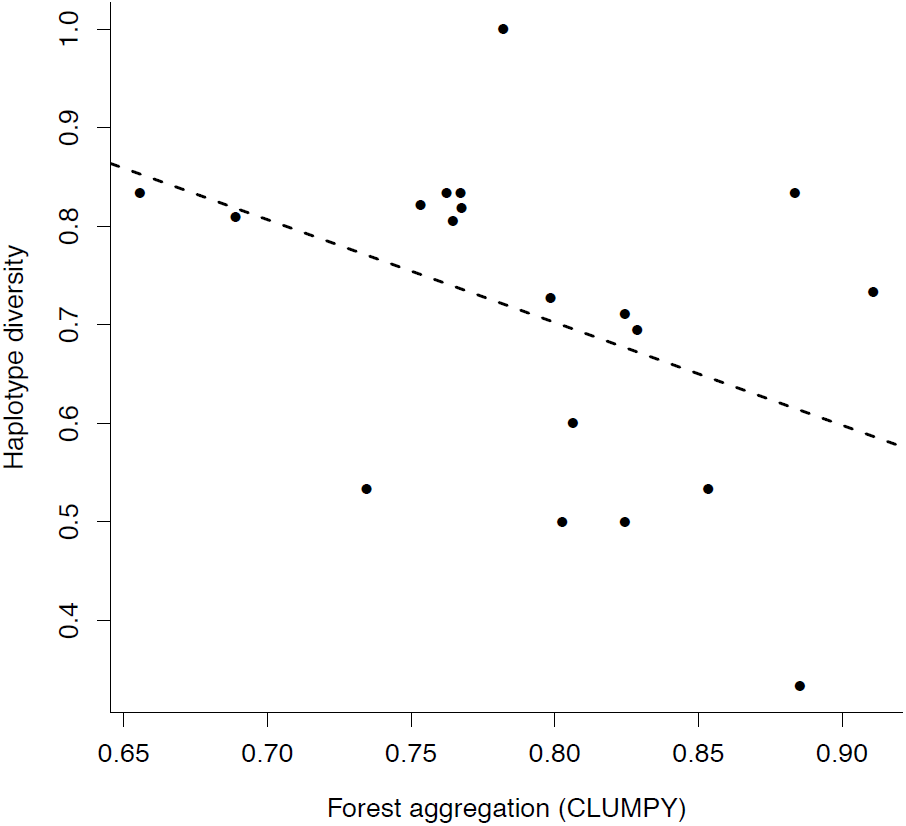
For *S. pygmaea*, haplotype diversity was lower in landscapes characterized by a higher degree of forest aggregation (CLUMPY).

### Spatial autocorrelation

We did not find any evidence of systematic patterns of spatial autocorrelation in the model residuals for any species (Fig. 5). After inspecting the spatial correlograms, we concluded that once the models incorporated the effects of forest configuration and composition, there were no broader spatial patterns in unexplained variation of within-population genetic diversity associated with geographic proximity.

**FIGURE 5.**
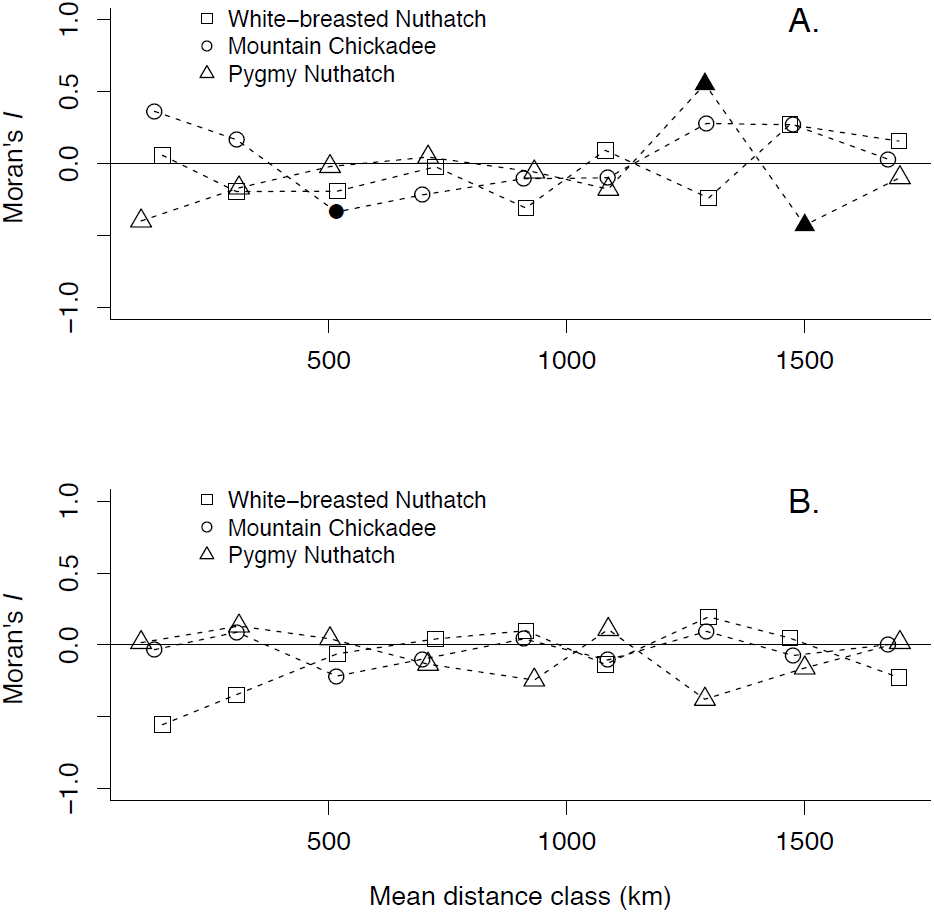
Spatial correlograms of GLM model residuals for *P. gambeli* (circles), *S. carolinensis* (squares) and *S. pygmaea* (triangles) for haplotype (A.) and nucleotide (B.) diversity. Filled symbols indicate significant spatial autocorrelation (*P* < 0.05) based on 10,000 permutations within a given distance class.

## DISCUSSION

We used existing genetic data on three forest breeding bird species to examine the relationship between within-population genetic diversity and forest fragmentation across multiple independent landscapes. We predicted that patterns of within-population genetic diversity with fragmentation would vary among species as a consequence of differences in their degree of habitat specialization or dispersal abilities. However, these patterns did not vary with life history traits in the directions that we predicted. Specifically, we predicted that within-population genetic diversity would be more likely to decline in response to habitat fragmentation among habitat specialists and among weak dispersers (Bailey et al. 2007) because they would be less likely to disperse through the intervening matrix between appropriate habitat fragments (and, therefore, would be more genetically isolated) than species with relatively broad habitat associations and stronger dispersal abilities. For two of the three species (*P. gambeli* and *S. carolinensis*), we found evidence that higher within-population genetic diversity was associated with landscapes characterized by lower levels of forest fragmentation. We found forest aggregation (CLUMPY) and the variability in forest patch area (AREA_CV) to be the two strongest landscape predictors of genetic diversity in these two species. These metrics captured the characteristics of habitat fragmentation, where large, compact clusters of forest with low variability in size supported populations with higher levels of within-population genetic diversity.

Our findings for *P. gambeli* and *S. carolinensis* are consistent with the predominant conclusion of a survey of studies comparing within-population genetic diversity between fragmented and control landscapes: in 26 of 45 comparisons (∼58%), studies found reduced genetic differentiation associated with fragmentation (Keyghobadi 2007). However, nearly half of these studies (*n* = 19) found no effect or even an effect in the opposite direction (Keyghobadi 2007). Likewise, in this study, we found patterns of higher within-population genetic diversity of *S. pygmaea* in fragmented landscapes. In our study, genetic diversity was higher in fragmented landscapes for the species that was the most specialized (Kingery and Ghalambor 2001, Spellman and Klicka 2006) and the most sedentary (Kingery and Ghalambor 2001), *S. pygmaea*. In contrast, *P. gambeli* and *S. carolinensis* have arguably broader habitat associations and stronger dispersal abilities than *S. pygmaea* (McCallum et al. 1999, Grubb and Pravosudov 2008), yet both showed a reduction of genetic diversity with fragmentation. Information about dispersal distances for these species (and for songbirds in general) is limited, and is likely to be greater than the distances reported in the literature (Tittler et al. 2009). Nevertheless, the patterns that we observed did not match our predictions based on their putative dispersal distances relative to one another, or with their relative specificity of habitat associations.

The elevated genetic diversity of *S. pygmaea* in fragmented landscapes is further surprising because of their cooperatively breeding social system. Many breeding pairs have one or more male helpers, usually offspring from pervious broods (Kingery and Ghalambor 2001). Some juveniles merge with neighbors from the previous winter’s group (Guntert et al. 1988, Sydeman et al. 1988). The short dispersal distances, natal philopatry, and habitat specificity of *S. pygmaea* are typical of many cooperatively breeding species (Woxvold et al. 2006, Haas et al. 2010). Such characteristics likely contribute to geographic isolation and genetic structuring among populations, and could increase the vulnerability of cooperative breeders to habitat loss and fragmentation (Koenig et al. 1996, Walters et al. 2004, Haas et al. 2010). However, cooperative breeders could be buffered from loss of genetic diversity in large fragments, from which juveniles might not have to disperse to find a breeding vacancy (Walters et al. 2004). More research is needed regarding how cooperative breeding could influence the relationship between fragmentation and genetic diversity.

Studying the relationship between habitat and within-population genetic diversity using contemporary landscape data can be problematic because variation quantified by molecular markers such as mitochondrial DNA may reflect historic, rather than contemporary, patterns of environmental variation (Frankham et al. 2010, Landguth et al. 2010, Sork and Waits 2010). As an example, *S. pygmaea* exhibited a positive correlation between genetic diversity and increased fragmentation and is the only species in the study with a single glacial refuge but a similarly large contemporary distribution (Spellman and Klicka 2006). By contrast, *P. gambeli* and *S. carolinensis* had lower diversity in fragmented landscapes and are phylogeographically structured into two or more reciprocally monophyletic clades and expanded to occupy their current distribution from two (or more) glacial refugia (Spellman and Klicka 2007, Spellman et al. 2007). Given the difference of range expansion from a single versus multiple glacial refugia, it is possible that the overall effective population size of *S. pygmaea* following expansion was larger and the contemporary distribution was achieved later than in *P. gambeli* or *S. carolinensis*. If this were true, then we would expect *S. pygmaea* to enter genetic drift equilibrium slower than other species with smaller, more disjunct postglacial population size. Thus, *S. pygmaea* may not have yet achieved this equilibrium, even though they have the lowest dispersal rates of the three examined species.

In some studies, when both historic and contemporary land use data have been studied, information about historic landscapes has proven to be a better predictor of current genetic variation than contemporary habitat fragmentation (Keyghobadi et al. 2005a, 2005b, Holzhauer et al. 2006). Mitochondrial markers generally reflect the genetic imprint of historical processes occurring hundreds of years before present (Frankham et al. 2010), but the samples used in this study were collected from natural areas less affected by modern day anthropogenic disturbance. Hence, it is likely that the contemporary patterns and variability of fragmentation across the study landscapes were sustained primarily by biophysical processes (e.g. soil type, topography, moisture gradients) rather than caused by human activity (e.g., human development). That being said, additional studies utilizing molecular markers (e.g., microsatellites) that may more readily capture contemporary genetic patterns could provide further insight into the relationships between patterns of landscape fragmentation and within-population genetic diversity we found.

Habitat fragmentation is a major driver of environmental change affecting avian populations across multiple levels of diversity (Fahrig 2003). In recent years, landscape genetics has focused on quantifying the effects of fragmentation and landscape connectivity on the genetic structure among populations (Luque et al. 2012), but studies must continue to take a landscape-scaled approach of quantifying how characteristics of habitat configuration are correlated with within-population genetic variation among a wide range of landscapes. Using genetic data from multiple populations across a broad geographic area, we found within-population genetic diversity was generally higher in landscapes characterized by less fragmented forest habitat for two out of three forest breeding birds. These findings support the prediction that fragmented landscapes sustain genetically less diverse populations; however, we found the opposite pattern for the species that would normally be considered most susceptible to fragmentation effects due to its limited dispersal abilities and habitat specificity. For broader generalization based on empirical data, studies on genetic diversity and multiple species of varying life-history traits inhabiting multiple landscapes are needed.

## ACKNOWLEDGMENTS

We thank the Carling/Benkman (University of Wyoming) and Dickinson/Dhondt (Cornell Lab of Ornithology) lab groups for constructive suggestions on an earlier version of this manuscript.

